# Genetic determinants of resistance and virulence among carbapenemase-producing *Klebsiella pneumoniae* from Sri Lanka

**DOI:** 10.1101/330951

**Authors:** Chendi Zhu, Veranja Liyanapathirana, Carmen Li, V. Pinto, Mamie Hui, Norman Lo, Kam-Tak Wong, N. Dissanayake, Margaret Ip

## Abstract

Whole genome sequencing of carbapenem-resistant Enterobacteriaceae from the intensive care units of a Sri Lankan teaching hospital revealed the presence of carbapenemase gene, *bla*_OXA-181_ among isolates of carbapenase-producing *Klebsiella pneumoniae* belonging to ST437 (2 strains) and ST147 (8 strains) in 2015. *bla*_OXA-181_ genes were carried in three variants of ColE-type plasmids. Elevated carbapemen resistance were observed in *ompK36* mutant strains. ESBL genes, plasmid–mediated quinolone resistance (PMQR) determinants (*qnr, aac(6’)-Ib-cr, oqxAB*) and mutations on chromosomal quinolone resistance-determining regions (QRDRs) with substitutions at ser83→I of *gyrA* and ser80→I of *parC* were observed. All strains possessed yersiniabactin on the mobile element ICE*kp* and other virulence determinants. Strict infection control and judicious use of antibiotics are warranted to prevent further spread of multidrug-resistant *Klebsiella pneumoniae* in the intensive care units.

## INTRODUCTION

Carbapenem-resistant *Enterobacteriaceae* (CRE) is a global threat. Among all the resistant mechanisms, plasmid-mediated horizontal carbapenemase gene transfer is the major acquired mechanism (1). Three types of carbapenemase (class A: *bla*_KPC_; class B: *bla*_NDM_, *bla*_IMP_, *bla*_VIM_; class D: *bla*_OXA-48-like_) can hydrolyze carbapenems at varying levels (2–4). High risk global clones such as ST258 *Klebsiella pneumoniae* are known to be associated with epidemic plasmids and these interactions are hypothesized to provide a survival advantage for these clones (5).

Sri Lanka is part of the Indian subcontinent. Although carbapenemases, like *bla*_NDM_ in nearby countries are well studied and spread worldwide, the single study currently available on CRE from Sri Lanka to date does not contain information on plasmids (6). Here we sequenced 10 CRE strains isolated from one Sri Lanka hospital using whole-genome sequencing (WGS) to describe antimicrobial resistant characteristics and the genetic profiles.

## RESULTS

### Resistant genes and virulence factors

During an eight-month period, all coliforms isolated from respiratory specimens from the intensive care units of a University Teaching Hospital in Sri Lanka were screened for carbapenem resistance using Stokes sensitivity testing method. Of these 64 single-patient isolates, 2.6% (10 isolates) were found to be resistant to carbapenems. Three of the strains were from the subsidiary ICU and 7 were from the main ICU. All 10 isolates were *Klebsiella pneumoniae* belonging to ST437 (2 strains) and ST147 (8 strains). Capsule information showed two *wzi* alleles (64 and 109). Their MIC data and antimicrobial resistant genes are listed in TABLE 1 and FIG 2. Only one carbapenemase gene, *bla*_OXA-181_ was found in all isolates. Extended-spectrum β-lactamase (ESBL) genes, including *bla*_CTX-M-15_, *bla*_SHV-11_, *bla*_TEM-1_ and *bla*_OXA-1_ were detected. Besides PMQR determinants (*qnr, aac(6’)-Ib-cr*, *oqxAB*), we also found the mutations on chromosomal QRDRs. Substitutions at ser83→I of *gyrA* and ser80→I of *parC* were observed in all strains, which have been frequently reported in quinolone-resistant *Klebsiella pneumoniae* worldwide (7). The virulence profile of 10 strains are the same, they all carry yersiniabactin genes (*ybt*, *irp1, irp2*, *fuyA*) and *kfu*, *mrk*. No *rmpA* or *rmpA2* gene were detected. Yersiniabactin genes were found on moble element ICE*kp.* 4437 core genomes and 2182 accessory genes were used to establish the pangenome dendrogram (see FIG 2). The accessory gene profile of ST437 and ST147 is quite different, making ST437 a distant cluster with others.

**TABLE 1.** Characteristics of the *bla*_OXA-181_ producing *K. pneumoniae* isolates

| Strain No. | Unit** | Stay length*<br>**<br>(days) | Specimen | ST | PlsmdSL | Other plasmids replicons | ompK36 PEFXG insertion | MICs(mg/L)* |  |  |  |  |  |  |  |  |  |  |
| --- | --- | --- | --- | --- | --- | --- | --- | --- | --- | --- | --- | --- | --- | --- | --- | --- | --- | --- |
|  |  |  |  |  |  |  |  | ETP | IPM | MEM | CTX | CAZ | AK | GN | CIP | TG | CT | FOS |
| SL33 | S | - | ET secretion | 437 | A | IncFIB, IncFII | No | 2 | 0.25 | 0.25 | >128 | 64 | 2 | 64 | 64 | 1 | 0.12 | 4 |
| SL49 | M | 7 | ET secretion | 437 | A | IncFIB | No | 4 | 0.5 | 0.25 | 128 | 128 | 4 | 0.5 | 128 | 1 | 0.25 | 4 |
| SL54 | S | - | ET secretion | 147 | C | IncFIB, IncR | No | 4 | 1 | 0.5 | >128 | 64 | 4 | 64 | 128 | 2 | 0.12 | 8 |
| SL34 | M | 4 | Sputum | 147 | B | IncFIB, IncA/C2, IncR | Yes | 32 | 4 | 32 | >128 | >128 | 4 | 64 | 128 | 0.5 | 0.5 | 8 |
| SL35 | M | 6 | Sputum | 147 | B | IncFIB, IncR | Yes | 32 | 4 | 32 | >128 | >128 | 4 | 128 | 64 | 0.5 | 0.25 | 16 |
| SL36 | M | 17 | Sputum | 147 | B | IncFIB, IncFII | Yes | 32 | 8 | 32 | >128 | 32 | 2 | 1 | 8 | 1 | 0.12 | 16 |
| SL46 | M | 9 | ET secretion | 147 | B | IncFIB, IncFII | Yes | 16 | 1 | 8 | 128 | 16 | 4 | 0.5 | 8 | 1 | 0.12 | 8 |
| SL50 | M | 13 | ET secretion | 147 | B | IncFIB, IncR | Yes | 32 | 8 | 16 | 4 | 0.5 | 1 | 128 | 32 | 1 | 0.25 | 32 |
| SL52 | S | - | ET secretion | 147 | B | IncFIB, IncR | Yes | 32 | 2 | 32 | >128 | >128 | 4 | 128 | 16 | 1 | 0.25 | 64 |
| SL68 | M | 4 | Sputum | 147 | B | IncFIB, IncR | Yes | 32 | 8 | 32 | >128 | >128 | 16 | 1 | 8 | 1 | 0.25 | 64 |

\* AK, amikacin; CAZ, ceftazidime; CIP, ciprofloxacin; CN, gentamicin; CT, colistin; CTX, cefotaxime; ETP, ertapenem; FOS,
fosfomycin; IPM, imipenem; MEM, meropenem; TG, tigecycline; FOS, fosfomycin.
\*\* M – Main study unit; S – Subsidiary units
\*\*\* Length of stay from date of specimen collection

**FIG 2.**
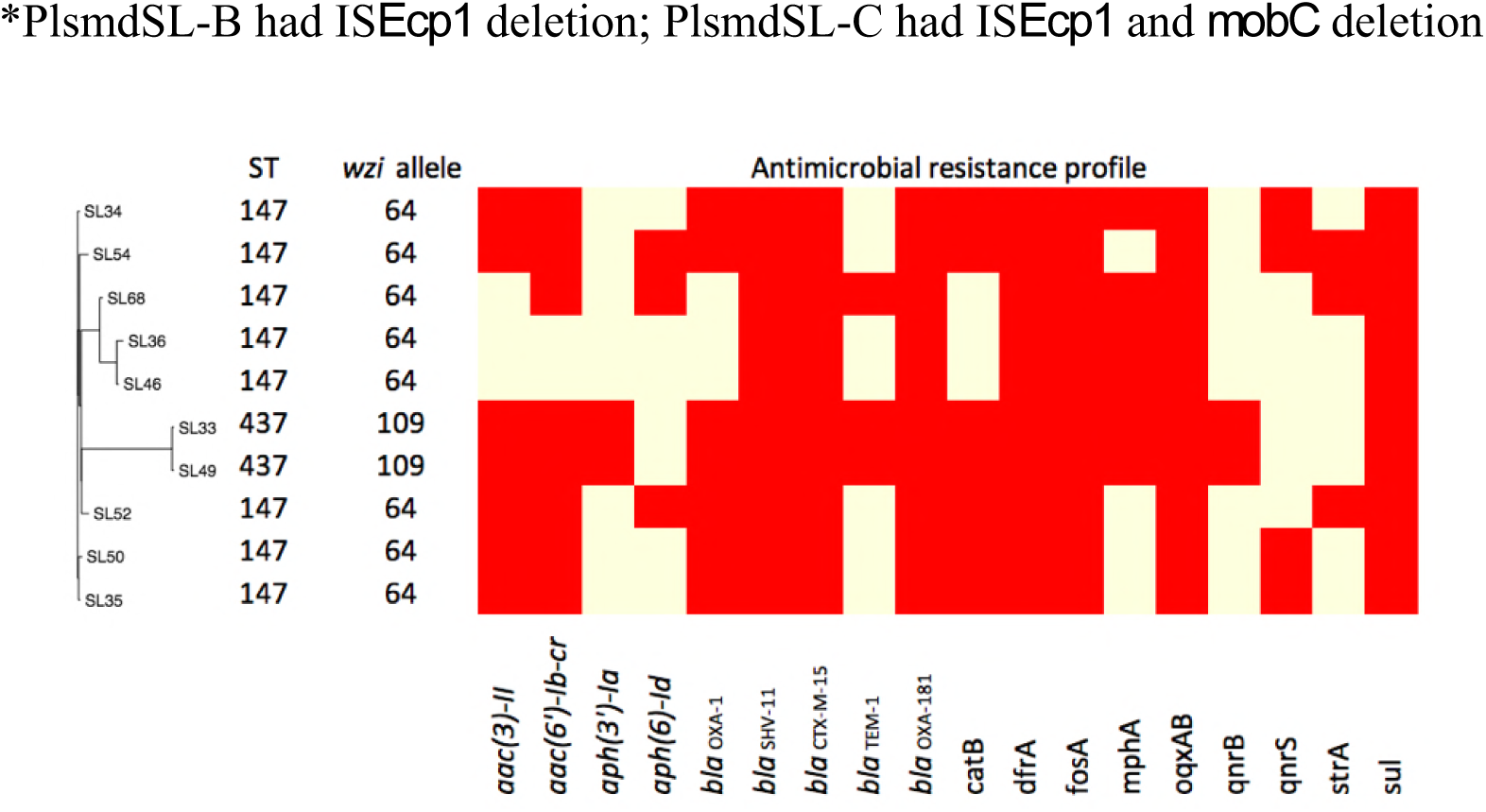
Pan-genome dendrogram of 10 Sri Lanka strains with annotation

### Elevated MICs of carbapenem

Carbapenem MICs of different strains varied. All the strains except SL33, SL49 and SL54 were resistant to ertapenem or meropenem at high level according to the CLSI guideline. In prior studies, mutation of *Klebsiella pneumoniae* outer membrane proteins *ompK35* and *ompK36* confer to increased MIC of carbapenems (2, 4, 8). We then aligned *ompK35* with wild-type one (GenBank accession no. AJ011501) and there were neither mutations nor insertions into *ompK35* and its promoter in all strains. *ompK36* was identical in six high resistant strains (*ompK36*-sl1). SL33, SL49 shared the same gene (*ompK36*-sl2), but SL54 owned an unique variation (*ompK36*-sl3). The difference of *ompK36*-sl1 and *ompK36*-sl2 was the insertion of Gly and Asp after PEFXG domain within the L3 loop and one mutation in loop L4 and alpha-helix respectively. PEFXG domain (porin size determinant) insertion was also observed in several studies contributed to the high resistant to carbapenems (2, 6, 8). *ompK36*-sl3 was different from others mainly between loop L3 and loop L6, but there was no insertion interruption in PEFXG domain.

### Plasmids harbored *bla*_OXA-181_

Three different ColE-type plasmids were identified (FIG 1). One (PlsmdSL-A) was identical to KP3 (GenBank accession no. JN205800) and was found in the two ST437 strains. PlsmdSL-A was a short plasmid with size of 7,606 bp and well described before (9). Another plasmid (PlsmdSL-B) had one insertion sequence (IS*Ecp1*) deletion compared with PlsmdSL-A. Several studies from USA, Germany, France have reported this plasmid in both *Klebsiella pneumoniae* and *Escherichia coli.* The third plasmid (PlsmdSL-C) had a mobile gene (*mobC*) deletion of PlsmdSL-B, which were not reported before.

**FIG 1.**
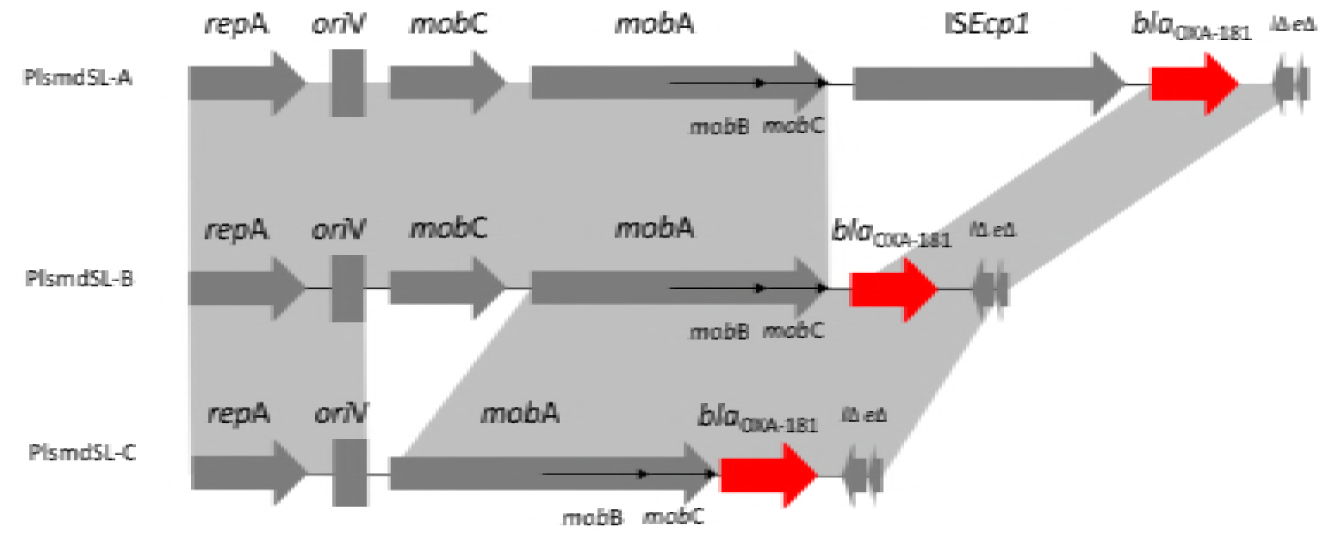
ColE-type plasmid

## DISCUSSION

Carbapenemase *bla*_OXA-181_ was first described in India as a Class D *bla*_OXA-48-like_ enzyme from clinical samples in year 2006 and 2007 (10). It is thought to be originated from an environmental strain as a chromosomal gene (11). Although it is detected worldwide, most of patients have a travel history to Indian subcontinent, especially India (3). In 2014, *bla*_OXA-181_ and *bla*_NDM_ was reported in *Klebsiella pneumoniae* in Sri Lanka of different STs, mainly ST14 and ST147 (6). ST437 is a single locus variant of the globally prevelant ST258, carrying *bla*_KPC_ in Brazil and *bla*_NDM_, *bla*_OXA-245_ (plasmid: IncL/M) in Spain (12). *bla*_OXA-181_ was previously described in ColE, IncT, IncX3 plasmids and chromosome, and all were isolated from patients transferred from India except IncX3 from China (9, 13–17). ColE plasmid encoded various β-lactamase gene and *bla*_OXA-181_ is related to transposon Tn*2013*. Insertion sequence IS*Ecp1* was considered related to *bla*_OXA-181_ acquisition, and its deletion may stabilize the resistant gene into plasmid (18). *mobC* gene deletion in current strain may affect the frequency of plasmid conjugal mobilization (19).

In this study, all strains harboured quinolone-resistant determinants with QRDR mutation on the chromosome. Recent epidemiology study has shown the correlation of quinolone consumption and CRE in US military health system, and another case-control study of CRE outbreaks in Netherlands have determined quinolone use as the only risk factor of acquisition of *bla*_OXA-48-like_ producing Enterobacteriaceae compared to other antibiotics use (20, 21). Possible mechanisms are co-transfer of two plasmids or recombination into one plasmid described before(16, 22).

Although all strains are nonhypervirulence *Klebsiella pneumoniae* (negative for *rmpA*/*rmpA2* genes, non K1/K2 capsule serotypes), they all encoded several yersiniabactin genes. These genes are on integrative conjugative elements (ICE*Kp*) can cause spread in *Klebsiella pneumoniae* population and as a predictor of invasive infection (23).

In conclusion, we discussed the genetic profile of multidrug resistant CRE in Sri Lanka. *bla*_OXA-181_ (through ColE-type plasmid) and yersiniabactin has disseminated to different STs of *Klebsiella pneumoniae.* We recommend active surveillance of high risk inpatients and long term studies to determine possible intra-unit transfer, especially as the length of stay in all instances where data was available is >72 hours. Judicious antibiotics use, especially quinolones is also recommended.

## MATERIALS AND METHODS

Single patient isolates were obtained from the respiratory specimens received from inpatients admitted to the intensive care units of the teaching hospitals of the University of Peradeniya between February to September 2015, During this period a total of 379 respiratory specimens were processed yielding 64 coliforms. Of these coliforms, 2.6% (10 isolates) were found to be resistant to carbapenems using Stokes sensitivity testing method. All of those isolates were saved for further study. The study was approved by the Institutional Ethical Committee of the Faculty of Medicine, University of Peradeniya.

Identification of all strains were confirmed by MALDI-TOF (matrix-assisted laser desorption ionization-time of flight mass spectrometry) at the Department of Microbiology, Chinese University of Hong Kong. The MICs of antimicrobials were determined by broth microdilution method according to Clinical & Laboratory Standards Institute guideline(CLSI) (24). Bacterial DNA was extracted with Wizard genomic DNA purification kit (Promega, Madison, USA). WGS was performed using the Illumina HiSeq 2500 platform, and unique index-tagged libraries were created for each sample to generate 90bp paired-end reads (Global Biologics, LLC).

The libraries gave 100x coverage for each strain on average. Quality control of the raw reads was performed by FastQC. Genomes were assembled using the SPAdes assembler (version V3.5.0) (25). Contigs of ≥ 500bp from each genome were included in the analyses. The Prokka (version 1.9) software was used for genome annotation, including ORF finding and gene function annotation (26). Raw reads and assembled contigs were used for multilocus sequence typing (MLST) analysis. SRST2 (Version 0.1.5) was used to map raw reads to the pubMLST database to infer ST, to the resistance gene database (ARG-ANNOT V3), to all the plasmid replicons in PlasmidFinder database(Updated 20170220) and to *K. pneumoniae* BIGSdb virulence gene database (http://bigsdb.web.pasteur.fr Accesstion date: 20180313) (27). The contig containing carbapenemase for each genome was blasted on NCBI to find the top hit plasmids (at least 95% identity and 95% coverage). Then we blasted the plasmids in our WGS data to extract possible plasmid contigs. Extracted contigs were further aligned to the reference plasmid using Mauve Alignment software to check similarity and coverage (28). Gaps were filled by PCR with primers designed based on sequencing data. Pan-genome dendrogram was created by Roary (29).

## ACKNOWLEDGEMENTS

We thank the Food and Health Bureau of Hong Kong Special Administrative Region for the support from Health and Medical Research Commissioned Fund.

